# Prefrontal-posterior coupling mediates emotion transitions and their influence on executive function

**DOI:** 10.1101/518860

**Authors:** Yu Hao, Lin Yao, Derek. M. Smith, Edward Sorel, Adam K. Anderson, Eric H. Schumacher, Gary W. Evans

## Abstract

Although emotions often result from dynamic experiences with self-regulation unfolding over time, most research has focused on responses to affective stimuli from a rather static perspective. We studied and analyzed emotion transitions, attempting to reveal brain functions related to affect dynamics. EEG responses were examined during exposure to stable versus changing emotion-eliciting images (static vs dynamic conditions) plus their impact on executive function (EF) assessed with the flanker task. During dynamic conditions, reduced prefrontal to posterior EEG coherence in the beta frequency band and greater left frontal activity occurred compared to the static conditions. Among individuals suffering higher chronic stress, subsequent EF was hindered after dynamic conditions. Furthermore, the adverse effects of emotion transitions on EF for more chronically stressed individuals were mediated by prefrontal-posterior coherence in low beta frequency band during emotional image sequences. Emotion appears to influence EF through changes in large-scale synchronization. Individuals high in chronic stress are vulnerable to these effects.

## Introduction

Emotional well-being is a key component of maintaining equanimity against a backdrop of variable experiences. Affect dynamics reflect self-regulation ability and are crucial for psychological health ^1–3^. Changing environmental conditions trigger affect dynamics, and studying the brain and behavior consequences provide insights about underlying mechanisms of cognition-emotion as they unfold over time ^4–7^. In our experimental paradigm, we define *emotion transition* as responses to *dynamic conditions*. As opposed to *static conditions* without affect transition, dynamic conditions are defined as exposure to stimuli of varying affective valence or intensity, e.g., image sequences with changing valence and/or intensity.

Cortical synchronization is related to a greater control of the affective state ^8^. Specifically, emotionally arousing/threatening stimuli and the modulation effect of executive control have been shown to influence EEG coherence between prefrontal and posterior cortical regions indexing changes in functional communication ^9, 10^. A reduction in the prefrontal– posterior coupling may reflect a loosening of top down control from the prefrontal cortex leading to greater emotional contagion ^9,11–15^. Executive function (EF) may also be recruited when conflicts are detected among different emotional affect between a current stimulus and reverberation from a previous emotional stimulus. Thus, emotional transitions may decrease prefrontal-posterior coherence.

The present study investigates neural responses to dynamic emotional conditions and their influence on subsequent EF as compared to static conditions. We also examine the influence of chronic stress on these outcomes. EF, emotion, and stress interact in an adaptive feedback loop in response to environmental cues ^16^. Individuals under chronic stress exhibit automatic responses that are more reactive than reflective, particularly in an unpredictable environment ^17, 18^, such as when presented with dynamic stimuli. Dynamic emotional stimuli can have cumulative effects which may be influenced by chronic stress. Although enhanced sensitivity to novel stimuli helps maintain response flexibility to new threats, it may add to the accumulative impact of glucocorticoids on the body potentially leading to mental illness ^19^. In addition, both acute and chronic stress can affect brain structures or functions ^20^, such as it influences prefrontal hemispheric imbalances in neuroendocrine regulation ^21^. Moreover, besides the deficits in regulatory processes of emotional salient events, the conflicts generated by affect transition recruit more cognitive control mechanisms in the prefrontal cortex ^22^. Processing dynamic emotional events likely requires more cognitive resources to regulate emotions and under chronic stress, these resources might be compromised. Therefore, dynamic stimulation might exhibit different aftereffects than purely negative events on EF.

In addition, lateralized activation of the prefrontal cortex is assessed because it has been linked to emotion-regulatory capacity ^23^. Our work links brain connectivity with responses to emotional transition events and executive functioning, and examines the moderating role of chronic stress in this relation. Thus, we hypothesized decreased coherence in dynamic conditions which, in turn, would mediate the relation between emotional stimulation and subsequent EF performance. Furthermore, we expected this link to be stronger among individuals experiencing elevated chronic stress because of inflexibility and/or diminished capacity to adapt to changes in the emotionally affective environment.

## Results

### Coherence in Static vs Dynamic Conditions

We examined beta frequency band [14 30] Hz, low [14 20] Hz and high [20-30] Hz beta frequency band coherence initially based on prior work, significant effects of condition on coherence were observed in both hemispheres. Because low beta frequency band ([14, 20] Hz) coherence had consistently strong effects and mediated the subsequent EF, we only report results from low beta frequency band coherence (Figure 1 (a-b). In the right hemisphere, *F (3, 96)* = 13.00, *p* < .001, proportion change of variance (*PCV_−within_*) = 26.7%; in the left hemisphere, *F (3, 96)* = 7.22, *p* < .001, *PCV_−within_* = 15.9%.

**Figure 1.**
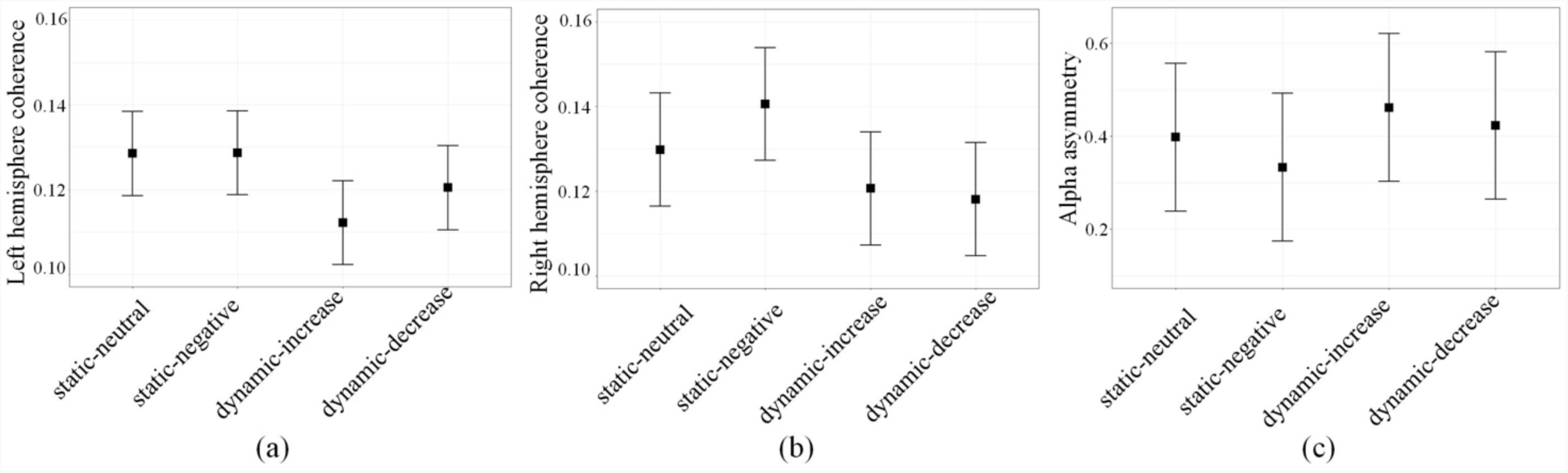
Low beta frequency band prefrontal-posterior coherence of four image sequence conditions in left (a) and right (b) hemispheres and frontal alpha asymmetry (c). Error bars are the standard errors.

Dynamic conditions evoked lower coherence than static conditions (Table 1). In the right hemisphere, *F (1, 98)* = 28.49, *p* < .001, *PCV_−within_ =* 21.7%; in the left hemisphere, *F (1, 98)* = 18.26, p < .001, *PCV_−within_ =* 14.9%. Dynamic conditions evoked greater left frontal activity than static conditions, *F (1, 98)* = 6.52, *p* = .01, *PCV_−Within_ =* 5.3% (Figure 2 (c)).

**Table 1.** Estimates (standard errors) of static vs dynamic condition

|  | Condition |  |  |
| --- | --- | --- | --- |
|  | Dynamic | Static | Dynamic-Static |
| Right hemisphere | 0.12 (0.01) | 0.14 (0.01) | -0.02 (0.00) |
| 95% CI | [0.10, 0.14] | [0.12, 0.16] |  |
| Left hemisphere | 0.12 (0.01) | 0.13 (0.01) | -0.01 (0.00) |
| 95% CI | [0.10, 0.14] | [0.11, 0.15] |  |
| Frontal alpha asymmetry | 0.44 (0.16) | 0.37 (0.16) | 0.08 (0.03) |
| 95% CI | [0.12, 0.76] | [0.06, 0.69] |  |

**Figure 2.**
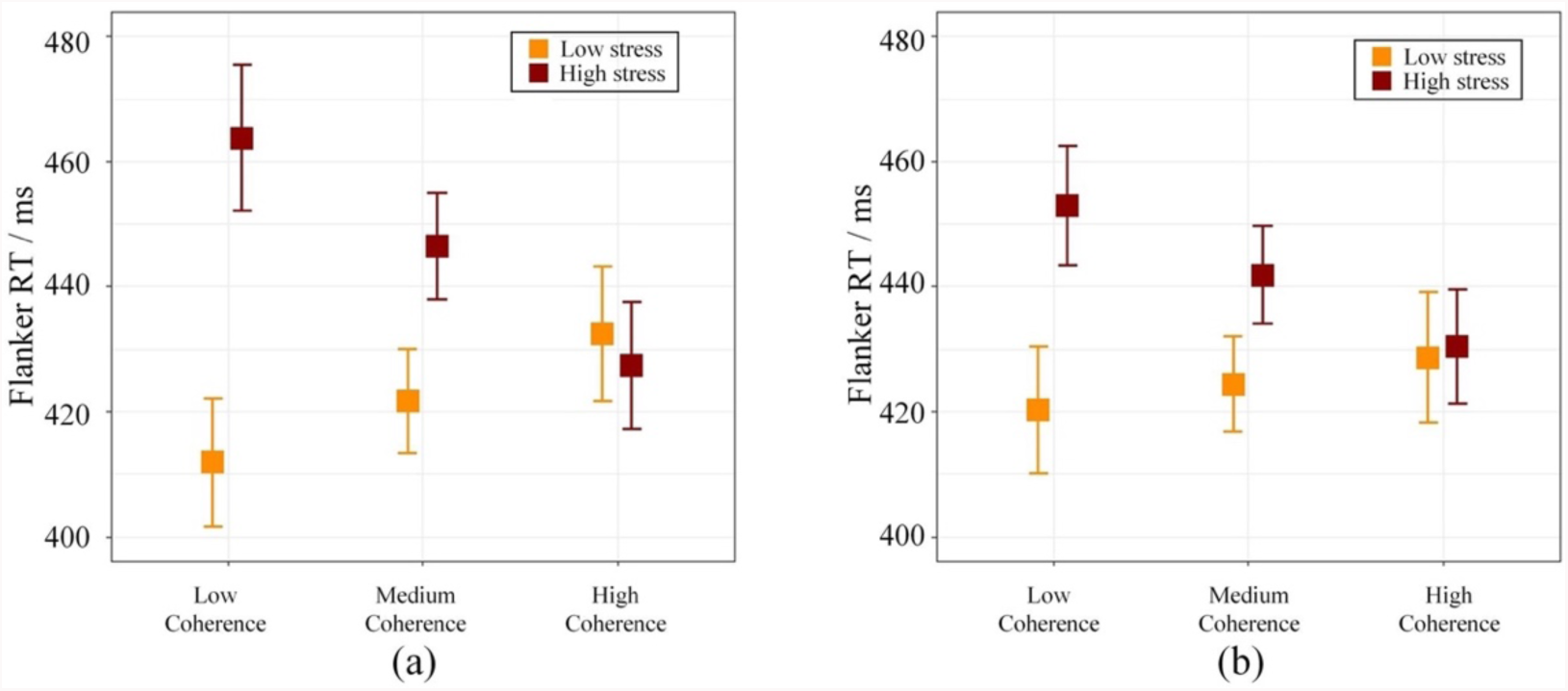
Flanker RTs by low beta frequency band coherence between prefrontal and posterior cortical regions during image sequence (a) and during flanker task (b) in the right hemisphere.

There were no main or interactive effects of chronic stress on coherence and frontal alpha asymmetry. Self-rated chronic stress (*M* = 15.91, *SD* = 6.33) was unrelated to gender, age, flanker performance or frontal-posterior coherence (with all tested frequency bands during image sequence and flanker task) and alpha asymmetry across conditions.

### Coherence as a Mediator of Condition Effects on Flanker

Chronic stress interacted with coherence during image viewing influence flanker RTs. In the right hemisphere, *F (1, 53)* = 17.70, *p* < .001, *PCV_−between_ =* 15.6%; in the left hemisphere, *F (1, 64)* = 10.99, *p* = .002, *PCV_−between_ =* 7.0%. Under high chronic stress, lower coherence during image sequences led to worse flanker performance; whereas higher coherence facilitated flanker performance, as depicted in Figure 2 (a).

The coherence during image sequence was highly correlated to coherence during the flanker task, *r* = 0.90, *t* (130) = 24.16, *p* < .001. Furthermore, similar trend as the influence of coherence in image sequence on flanker RTs, the influence of coherence during flanker task on RTs was dependent on chronic stress in the right hemisphere, *F* (1, 68) = 4.89, *p* = .030, *PCV_−between_ =* 16.9%. Under high chronic stress, lower coherence during flanker task led to worse flanker performance; whereas higher coherence facilitated flanker performance, as depicted in Figure 2 (b).

The effect of static vs. dynamic conditions on flanker performance was mediated by coherence during image sequences, only in participants suffering high chronic stress (above the mean) (Figure 3). In the right hemisphere, the indirect path *ab* = 7.91, *standard error* = 3.20, *p* = .013; in the left hemisphere, the indirect path *ab* = 5.01, *standard error* = 2.48, *p* = .044. The dynamic conditions evoked lower coherence that resulted in slower RTs among those with high chronic stress. Among those emotion or attention related EEG (i.e., midline theta band power, theta/beta ratio, high beta/low beta ratio, and alpha asymmetry), only anterior-posterior low beta coherence during emotional conditions appeared to affect RTs.

**Figure 3.**
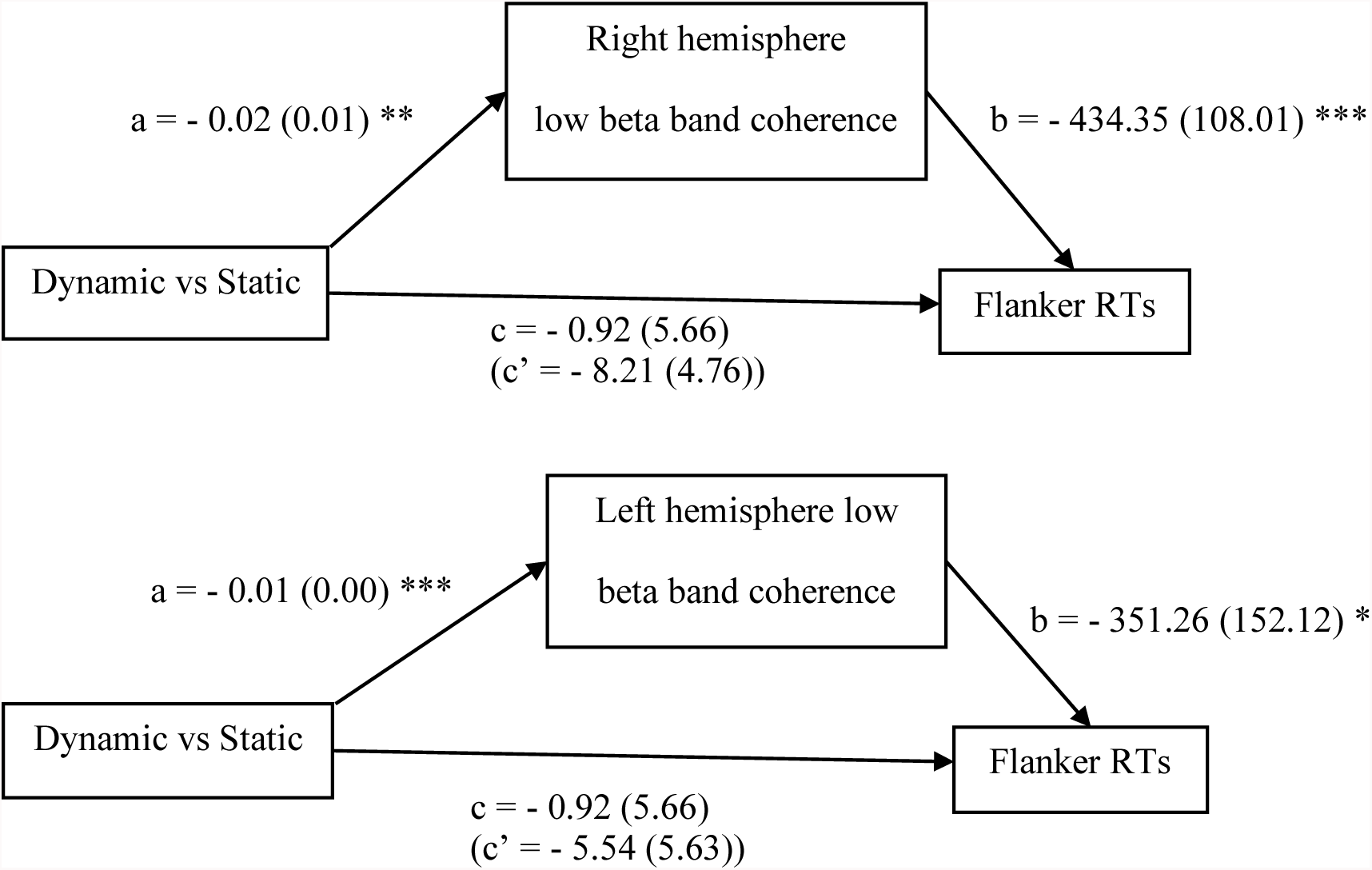
Mediation diagrams—prefrontal to posterior EEG coherence in the low beta band frequency band mediates the effects of emotion stimulation conditions on flanker RTs. Estimate (standard error), * p <0.05, ** p <0.01, *** p<0.001.

## Discussion

Prefrontal-posterior EEG coherence in beta frequency band coherence is lower during emotion transition regardless of affect (neutral or negative). Negative stimuli influence the beta frequency band coherence ^10^, but this is the first demonstration that beta frequency band coherence between prefrontal and posterior cortical regions is related to dynamic changes in emotion—*emotion transition*. The coherence in low beta ([14-20] Hz) frequency band also mediated emotional stimuli on subsequent EF among chronically stressed individuals.

In our experimental paradigm, we consider the interaction of changes in emotional stimuli and neural responses thereto as potential correlates of self-regulation. Decreases in beta frequency band prefrontal-posterior EEG coherence induced by emotional stimulation reflect a loosening of the prefrontal cortex’s regulatory control of parietal regions ^9,11–15^. Therefore, beta coherence plays a critical role in brain networks responding to emotion-cognition interaction. Image sequences with emotion transitions invoke more regulation effort as left frontal activity increases in comparison to responses to static neutral or negative images ^23^.

Our study calls attention to the mediating role of brain synchronization in the relationship between the affective dynamism of stimulation and EF. In the present study, EF in chronically stressed individuals is hindered when preceded by reduced prefrontal-posterior neural uncoupling under emotional transitions. Residual emotion from preceding images might generate emotional conflicts, impelling more executive control mechanisms for self-regulation to manage conflicting emotional stimuli. To having greater conflict, these individuals might be less able to manage conflicts ^24^. This also appears to diminish at least short term EF as indexed herein by the flanker task ^25^. With the reduced coherence evoked by emotion transitions continued during the flanker task, chronically stressed individuals were more vulnerable and showed deficits in inhibitory ability during flanker task. This is salient among persons suffering greater chronic stress perhaps because of already constrained self-regulation capacity.

The primary limitations of the current study are that the duration of the induced drop in EF performance is unknown because only 6 trials of flanker task manifested aftereffect. We also do not know if other negative emotional sequences would create similar results. A valuable extension of this study would be to examine persons with self-regulatory difficulties, for instance, individuals with anxiety disorders. Reversely, manipulating self-regulation capability by EF training ^25, 26^ or cognitive reappraisal ^27^ under emotion transition and its research on psychopathology are needed. Moreover, the investigation of emotion transition can be extended into other forms of dynamics, such as including more valence changes, or multiple transitions in affect stimulations. Limitations of the method (magnitude-squared coherence) include possible volume conduction artifacts. However, the relative difference of coherence among different experimental conditions is of our primary interest. In our study, no significant changes occurred in delta or theta EEG coherence comparing dynamic vs static condition during affective viewing. This also argues against general volume conduction confounds.

Emotion transitions may have more significant effects on coupling in neuronal regions and EF measures than static events without transitional affect. Brain responses to dynamic stimuli may reflect emotional regulation capacity specifically and coping skills more generally.

## Methods

### Participants

Participants were 40 right-handed students from Cornell University. Informed consent was obtained, and participation was compensated with course credits or $20 cash. Due to incomplete data, 33 participants were included in the final sample (51.5% females, age M = 22.40, SD = 3.80). Exclusion criteria were any open or healing wounds on the scalp, use of any medication that could affect nervous system processing and any history of neurological disorders. Participants were requested to refrain from alcohol caffeine, and other stimulants for four hours prior to the experiment. They were also asked to sleep for at least six hours the night before the experiment. The study was approved by Cornell’s Institutional Review Board and was performed in accordance with the its guidelines. Informed consent was obtained from all participants.

### Procedure

To measure chronic stress, we administered the Perceived Stress Scale (PSS) before EEG recordings ^28^. PSS was designed to measure perceived stress level, i.e., unpredictable, uncontrollable, and overloaded. This questionnaire contains 10 items and requires respondents to rate the degree to which situations are appraised as stressful in the last month, such as: “In the last month, how often have you felt confident about your ability to handle your personal problems?” or “In the last month, how often have you felt difficulties were piling up so high that you could not overcome them?”. PSS is a reliable and valid instrument for assessment of perceived stress in college students and workers ^29, 30^.

Affective state was manipulated by 96 negative, emotionally threatening images (valence: *M* = 2.37, *SD* = 0.65, arousal: *M* = 5.95, *SD* = 0.77) and 96 low arousal neutral images (valence: *M* = 5.12, *SD* = 0.53, arousal: *M* = 3.17, *SD* = 0.66) selected from IAPS database ^31^. Four conditions of *image sequences* were randomly presented with no image repetition to participants. For dynamic conditions, the images either transitioned from neutral to negative (dynamic-increase) or from negative to neutral (dynamic-decrease), which appeared 18 times each. For static conditions, the sequences consisted of all neutral (static-neutral) or all negative (static-negative) images, which appeared 6 times each. Each sequence involved four images with each image 4500 ms duration. First two images and last two images were from the same affect category, therefore, the transition occurred at the third image. More dynamic image sequences ensured the experiment conditions were in a dynamic setting. To counteract the balance of the amount of image sequence between static and dynamic conditions, we selected 6 sequences in dynamic conditions that were next to the static conditions. These were distributed across the experiment.

Immediately after each sequence, 12 trials of the flanker task ^32^ each 1200 ms (stimulus 200 ms and response window: 900, 1000 or 1100 ms) were presented to assess EF performance ^33^. In the flanker task, a visual array of flag stimuli with the middle flag randomly surrounded by flags in the same direction (congruent) or in the opposite direction (incongruent). Participants responded to the direction of the middle flag as quickly and accurately as possible.

### EEG Processing

EEG was recorded from a 128-channel BioSemi EEG device with digital sampling rate at 512 Hz. Coherence measures the degree of covariance between two spatially distinct signals in prefrontal and posterior regions. We applied the magnitude-squared coherence, which was calculated by the cross-spectrum divided by the product of the auto-spectrum of the two signals. It includes information of the amplitude and phase. The phase difference between the two signals plays an important role in estimating brain connectivity ^34^. Following the methods described in Miskovic and Schmidt ^10^ and Allen et al. ^35^, the coherence in beta frequency band [14 30] Hz, low beta frequency band [14 20] Hz and high beta frequency band [20-30] Hz and frontal alpha asymmetry at the dorsolateral region were calculated throughout the image sequence (18000 ms) for each condition. For the full description of the EEG processing, see supplement. We followed up these hypothesized effects with exploratory analyses of other EEG metrics that are related to emotion and attention.

### Statistical analysis

We applied linear mixed effect models to predict brain activities by image sequence condition with each hemisphere and each frequency band. Then we added chronic stress as interaction with the condition. High and low chronic stress are plotted ±1 SD from the mean for descriptive purposes only in the Figures. Inferential analyses maintained the continuous nature of the chronic stress variable.

To explore condition effect on subsequent EF, and the mediation role of brain activities, we first predicted flanker reaction times (RTs) by condition. Then RTs was predicted by brain activities under image sequence and flanker task. Again, chronic stress was added as an interaction term.

Flanker performance was calculated as RTs on incongruent trials with RTs on congruent trials as a covariate. Trials with inaccurate responses or outlier RTs were deleted (3.14% of trials were deleted). We found that the first half of flanker trials (6 trials) followed by each image sequence had stronger interaction effect with chronic stress while the effect decayed in the end given temporal separation from the emotional stimuli. Thus, we used the initial six flanker trials following the emotional stimuli in the following analysis.

## Acknowledgements

We are grateful to the participants. We thank James Rounds and Qiuyan Sun for their assistance with data collection. We also thank Cornell Statistical Consulting Unit for statistical analysis advice. This work is supported by the Orrilla Wright Butts, Home Economics Extension, Jean Failing and Virginia F. Cutler Fellowships to Y. Hao.

## Contributions

Y.H. and G.W.E. developed the study concept. Y.H contributed to the study design. Data collection was performed by Y.H. Y.H and L.Y. performed the data analysis. Y.H and D.M.S. performed the data interpretation. Y.H. drafted the manuscript, and L.Y., D.M.S., E.S., A.K.A., E.H.S. and G.W.E. provided critical feedback.

**Competing financial interests**
The authors declare no competing interests as defined by Nature Research, or other interests that might be perceived to influence the results and/or discussion reported in this paper.

## Data availability

The datasets generated during and/or analysed during the current study are available from the corresponding author on reasonable request.

## Supplement

### EEG signal processing

EEG was recorded from a 128-channel BioSemi EEG device with digital sampling rate at 512 Hz. All EEG channels were referenced offline to the algebraic average of left and right mastoids and notch filtered (55~65 Hz) to remove power-line noise. EEG signals were bandpass filtered between 1~40 Hz. Bad channels were identified and spherically interpolated. Then the data were epoched from 4 seconds before and 35 seconds (total image sequence and flanker task duration) after the sequence onset. Each epoch was visually inspected. Those epochs with obvious abnormal signal segments, such as head movement, were excluded for Independent Component Analysis (ICA). Then ICA (EEGLab toolbox ICA function) was utilized to detect and remove artifact contaminated by eye movements, muscle, and cardiac artifacts. After removing the artifact components, the ICA source signals were transferred back to the original signal space, which was then used for the subsequent analysis.

The first half second of affective image viewing (0 to 0.5 s) was removed from analyses to eliminate sensory transition effect. The average number of artifact free seconds per participant was *M* = 79.90 (*SD* = 4.61), *M* = 79.80 (*SD* = 4.39), *M* = 238.4 (*SD* = 11.80) and *M* = 239.2 (*SD* = 9.20) for conditions of static-neutral, static-negative, dynamic-increase and dynamic-decrease respectively. There was no significant difference in the time length between static conditions (p = .910), and between dynamic conditions (p = .720).

Artifact-free EEG data were then submitted to a discrete Fourier transform with a Hamming window of 2000 ms width and 50% overlap, and Welch's method was used to estimate the auto spectrum of itself and cross spectrum between two signals. Electrodes distance is a factor that influences the effect of volume conduction. The spatial resolution of EEG is approximately 5 cm ^36^ and the optimal distance between electrodes must be around 10–20 cm in human EEG-recordings to minimize the effect of volume conduction ^37, 38^. We confined our analyses to electrode pairs located no less than ~ 18 cm from each other to minimize the effect of volume condition. Following Miskovic and Schmidt (2010), four clusters of electrodes were selected, with right frontal C16, C10, C7; left frontal C29, C32, D7; right parietal B4, B11, A28; left parietal A7, D31, A15. Coherence scores of nine electrode pairs each were averaged to summarize interaction within the different brain region, respectively (right hemisphere: C16-B4, C16-B11, C16-A28, C10-B4, C10-B11, C10-A28, C7-B4, C7-B11, C7-A28; left hemisphere: C29-A7, C29-D31, C29-A15, C32-A7, C32-D31, C32-A15, D7-A7, D7-D31, D7-A15).

The cross-spectral coherence between two channels was calculated using the following formula in (1). (Note: *S*_*xy*_ denotes the cross-spectrum, *S*_*x*_ and *S*_*y*_ denotes the auto-spectrum; The E denotes the expectation across the repeated sequences).

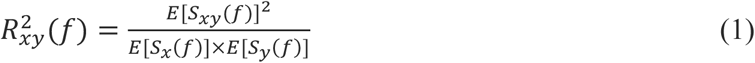

Alpha asymmetry was calculated following Allen et al. (2004). The auto spectrum of the frontal channels (i.e., F3 and F4 in 10-20 system, C32 and C10 in BioSemi system) was calculated using the Welch’s method with a hamming window of 2000 ms width and 50% overlap. Then the power within alpha frequency band ([8 13] Hz) is summed up to extract the alpha power. The alpha asymmetry is defined as ln(*P_F4_*)-In(*P_F3_*). To have a reliable task-related estimation, the power within each condition of image sequences is averaged to estimate the corresponding task-related power and its alpha asymmetry.

